# Rapid and Repeated Local Adaptation to Climate in an Invasive Plant

**DOI:** 10.1101/420752

**Authors:** Lotte A. van Boheemen, Daniel Z. Atwater, Kathryn A. Hodgins

## Abstract

- Biological invasions provide opportunities to study evolutionary processes occurring over contemporary timescales. To explore the speed and repeatability of adaptation, we examined the divergence of life-history traits to climate, using latitude as a proxy, in the native North American and introduced European and Australian ranges of the annual plant *Ambrosia artemisiifolia*.
- We explored niche changes following introductions using climate niche dynamic models. In a common garden, we examined trait divergence by growing seeds collected across three ranges with highly distinct demographic histories. Heterozygosity-fitness associations were used to explore the effect of invasion history on potential success. We accounted for non-adaptive population differentiation using 11,598 SNPs.
- We revealed a centroid shift to warmer, wetter climates in the introduced ranges. We identified repeated latitudinal divergence in life-history traits, with European and Australian populations positioned at either end of the native clines.
- Our data indicate rapid and repeated adaptation to local climates despite the recent introductions and a bottleneck limiting genetic variation in Australia. Centroid shifts in the introduced ranges suggest adaptation to more productive environments, potentially contributing to trait divergence between the ranges.

## INTRODUCTION

During biological invasions species commonly spread over large and climatically diverse geographic areas. In doing so, they often re-establish within climatic niches found in their native ranges, and in some cases flourish in new environments (Sax & Brown, 2000; Allendorf & Lundquist, 2003; Atwater *et al.*, 2018). Although plasticity and broad ecological tolerance may facilitate the spread of invaders across such heterogeneous climates (e.g. Geng *et al.*, 2007; Zhang *et al.*, 2010), a growing number of empirical examples suggest that rapid adaptation to local conditions can also enable the establishment and spread of these species in the face of new selective dynamics (Huey *et al.*, 2000; Lachmuth *et al.*, 2011; Colautti & Barrett, 2013; Chown *et al.*, 2014; Turner *et al.*, 2014; Oduor *et al.*, 2016). As such, invasions provide an opportunity to study contemporary adaptive processes, which is key in an era of rapid, human-induced, environmental change. As many single species have invaded several distinct regions of the globe, comparisons of the native range to multiple successful introductions could illuminate if and when traits evolve in parallel along climatic gradients (Moran & Alexander, 2014).

Climate is known to be an important selective factor shaping a diverse array of plant traits from physiological (e.g. Maron *et al.*, 2007; Ordoñez *et al.*, 2009), to life history traits (e.g. Franks *et al.*, 2007; Nakazato *et al.*, 2008; Colautti & Barrett, 2013) to defence (e.g. Moles *et al.*, 2011; Colomer-Ventura *et al.*, 2015). For instance, specific leaf area commonly increases with latitude (Frenne *et al.*, 2013), likely reflecting adaptation to latitudinal changes in temperature, precipitation, light availability, or herbivory (Poorter *et al.*, 2009). Furthermore, trade-offs among life history traits can shape adaptive trait divergence in response to local conditions and impact the evolutionary trajectory of trait combinations in invasive populations (Etterson & Shaw, 2001; Griffith & Watson, 2005; Colautti *et al.*, 2010; Hodgins & Rieseberg, 2011; Colautti & Barrett, 2013). Reductions in season-length at higher latitudes are frequently reported to select for early flowering at the cost of diminished plant size (Colautti *et al.*, 2010; Li *et al.*, 2014). Coordinated shifts in life-history traits along latitudinal gradients within ranges have been documented in several invasive plants (Dlugosch & Parker, 2008b; Hodgins & Rieseberg, 2011; Colautti & Barrett, 2013).

Latitudinal patterns in plant size could have important evolutionary consequences for other plant traits. Variation in plant size can influence optimal resource allocation to male and female sex function (Charnov, 1982; De Jong & Klinkhamer, 1989; Klinkhamer *et al.*, 1997). In wind-pollinated plants, height can affect fitness returns directly through more effective pollen dispersal in taller plants (Burd & Allen, 1988; Klinkhamer *et al.*, 1997), as well as indirectly through increased availability of resources (Lloyd & Bawa, 1984; De Jong & Klinkhamer, 1989; Klinkhamer *et al.*, 1997; Zhang, 2006). Outcrossing wind-pollinated plants are predicted to adaptively change sex allocation to be more male-biased with increase in size (Lloyd, 1984; De Jong & Klinkhamer, 1989; de Jong & Klinkhamer, 1994; Klinkhamer *et al.*, 1997), as local seed dispersal should lead to saturating female gain curves (Lloyd & Bawa, 1984; Sakai & Sakai, 2003), yet more linear male function gain curves are expected (Klinkhamer *et al.*, 1997; Friedman & Barrett, 2009). Latitudinal clines in height could therefore be expected to lead to adaptive shifts in sex allocation. However, studies investigating the evolution of sex allocation patterns over wide geographic ranges are rare (Guo *et al.*, 2010; Barrett & Hough, 2012).

Many factors could impact trait evolution of native and invasive populations evolving in response to similar climatic gradients resulting in divergent outcomes. For instance, demographic events, such as bottlenecks, genetic drift in founding populations and admixture could differentially affect native and invasives’ adaptive capacity or influence the route by which adaptive evolution proceeds (Lee, 2002; Facon *et al.*, 2006; Prentis *et al.*, 2008; Rius & Darling, 2014; Bock *et al.*, 2015; Estoup *et al.*, 2016; Hodgins *et al.*, 2018). Although the impacts of bottlenecks during colonization on molecular variation are well characterized its effect on quantitative trait variation are not well established (Dlugosch & Parker, 2008a; Dlugosch *et al.*, 2015a). Moreover, even if the required genetic variation is present in the introduced range, the probability of observing trait clines in the introduced range depends on the time since introduction and the strength of climate-mediated selection (Prentis *et al.*, 2008; Bock *et al.*, 2015). Shifts in the biotic environment associated with introduction may also influence the evolution of plant traits, potentially impacting trait clines. Indeed, the impact of the biotic environment has been commonly considered in the examination of trait evolution in invasive plant species (Felker□Quinn *et al.*, 2013). By contrast, the invasion history could create patterns of trait variation (e.g., through climate matching (Maron *et al.*, 2004)) mimicking adaptive population divergence (Colautti *et al.*, 2009; Colautti & Lau, 2015). Therefore careful consideration of the source populations is required during studies examining adaptive trait evolution during invasion. Dissection of these various mechanisms is required to advance our understanding of the role of rapid evolution in invasive spread (Keller *et al.*, 2009; Bonhomme *et al.*, 2010; Lachmuth *et al.*, 2011; Agrawal *et al.*, 2015; Cristescu, 2015; Dlugosch *et al.*, 2015a).

We examine the repeatability and divergence of important life-history traits in the native North American and introduced European and Australian ranges of *Ambrosia artemisiifolia*. We raised seeds collected at 77 locations from broad climatic scales in a common garden and accounted for non-adaptive genetic differentiation using 11,598 genotype-by-sequencing SNPs, as neutral processes could impact trait variation. We here focus on evolution of plant size (height, biomass and growth), reproductive traits (flowering onset, dichogamy, sex function allocation, seed weight, total and relative reproductive biomass) and physiology (specific leaf area). We investigate four specific questions: *1) Do native and introduced populations occur in similar climates niches?* As climate is likely important in governing trait variation in this species, we examined climatic niche shifts following introduction to assess how traits might be predicted to diverge within and among the ranges. *2) Is there evidence for rapid parallel adaptation to latitude?* Repeatable trait clines for each range along latitudinal gradients, highly associated with many aspects of climate, would provide strong evidence that rapid adaptation to similar selective environments has occurred. We additionally explore coordinated shifts in traits potentially linked by trade-offs. *3) Is there evidence for trait differentiation between native and introduced ranges?* By examining patterns across multiple introduced ranges, we can explore if novel recipient communities generated trait divergence during introduction, or if adaptation to local climates dominates patterns of trait variation. *4) Is there a correlation between heterozygosity and fitness related traits?* Significant correlations would reveal if demographic changes such as bottlenecks, admixture and inbreeding have likely impacted the evolution of traits during this species’ extensive range expansion. Such correlations are predicted at the individual and population level in regions that have recently expanded their range, including those that have undergone admixture (Peischl & Excoffier, 2015).

## MATERIAL AND METHODS

### Study species

*Ambrosia artemisiifolia* is a wind pollinated, outcrossing, hermaphrodictic annual, which has aggressively spread from its native North America to many regions worldwide (Laaidi *et al.*, 2003; Smith *et al.*, 2013). The first records documenting the invasion are in central France around 1850 (Chauvel *et al.*, 2006). Multiple introductions from distinct native sources ensued to both east and west Europe, resulting in levels of genetic variation equivalent to those found in the native range (Chun *et al.*, 2010; Gladieux *et al.*, 2010; Gaudeul *et al.*, 2011; van Boheemen *et al.*, 2017). Genetic analysis suggests the Australian populations originate from a subsequent single introduction event around 80 years ago, derived from the European introduction, although the exact source is unknown (van Boheemen *et al.*, 2017). Range expansion likely occurred both north- and southward following this south-Queensland introduction (Palmer & McFadyen, 2012; van Boheemen *et al.*, 2017)

Latitudinal clines in phenology have been observed within the native North American range and the introduced ranges of Europe (Chun *et al.*, 2011; Leiblein-Wild & Tackenberg, 2014) and China (Li *et al.*, 2014), with earlier reproduction and greater relative investment in reproductive biomass in high-latitudinal compared to low-latitudinal populations (Chun *et al.*, 2011; Hodgins & Rieseberg, 2011; Li *et al.*, 2014). The wind-spread pollen is a major cause of hayfever worldwide and its medical treatment costs millions of dollars each year (Taramarcaz *et al.*, 2005), providing considerable incentive to understand the factors impacting pollen production in this species.

### Climate niche dynamics

To estimate the climatic niche occupied by *A. artemisiifolia* in its native North American, introduced Eurasian and Australasian ranges, we used ordination-based species distribution models. Models were taken from a larger study of 835 species (Atwater *et al.*, 2018), where the methods are described in detail. Briefly, occurrence data were collected from the Global Biodiversity Information Facility and plotted in 2-dimensional climate space based on rotated component variables (RCA1: temperature and RCA2: precipitation) of the 19 WorldClim variables (Hijmans *et al.*, 2005) representing annual trends, seasonality and means in temperature or precipitation (Table S1, Supplementary Material). We selected these variables as they are commonly used in studies on species distribution and local adaptation including for studies of *A. artemisiifolia* (e.g. Leiblein-Wild & Tackenberg, 2014; Sun *et al.*, 2017). We next generated a climatic occupancy map that described the occurrence probability of *A. artemisiifolia* as it varied among climates. Next, we divided this occupancy map by a map of sampling bias, as estimated from a dataset of 815 terrestrial plant species (Atwater *et al.*, 2018). Removal of both geographic and climatic sampling bias in this way produced bias-corrected estimates of occurrence probability in each set of climatic variables, for each of the three ranges (North American, Eurasian, and Australasian). Finally, these predictions were smoothed using We used Monte Carlo resampling (*n* = 120) to compare observed niche dynamics to those expected using randomly resampled occurrence points (Atwater *et al.*, 2018). We used Schoener’s *D* (Schoener, 1968) to estimate niche overlap between the native range and each introduced range (*i.e.* the degree of similarity in occupancy probabilities between both ranges), and we estimated niche stability (the proportion of climates occupied by the species in both their introduced an native range), expansion (the proportion of climates occupied in the introduced range that are available but unoccupied in the native range), and unfilling (the proportion of climates occupied in the native range that are available but unoccupied in the introduced range)(Guisan *et al.*, 2014). Finally, we tested whether the location of the niche centroid differed between native and introduced ranges.

### Data collection

To investigate local environments, we described climatic differences between 27 populations in the native range of North America, 32 populations in the introduced European range, and 18 populations in the introduced Australian range (Fig. S1, Supplementary Material). We used the 19 WorldClim variables and added a geographic dimension to the data by including altitude, latitude and longitude, as these variables are shown to be important in *A. artemisiifolia* growth and fitness (Chun *et al.*, 2011; Chapman *et al.*, 2014). To explore associations between climatic and geographic variables in the sampled populations, we applied a principal component analysis (PCA). We opted to present latitudinal trait clines only, as *i)* the primary principle component was highly correlated with latitude (Fig. S2); *ii)* precipitation and temperature variables were strongly associated with latitude (Spearman’s ρ^2^=0.326-0.417 (Table S2); *iii)* exploratory analyses revealed associations between each trait and many climatic and geographic variable were highly similar to trait~latitude trends (results not shown); and *iv*) is shown to affect *A. artemisiifolia* season length and photoperiod (Ziska *et al.*, 2011).

To assess the potential for adaptive differentiation along latitudinal gradients, we measured trait variation in a common garden of raised seeds collected at broad geographical scales across the three ranges. This is a traditional approach to detect genetic differences among populations (e.g. Bossdorf *et al.*, 2005; Colautti *et al.*, 2009; Hodgins & Rieseberg, 2011; Savolainen *et al.*, 2013; de Villemereuil *et al.*, 2016). We collected seeds in 2013-2014 and randomly selected an average of 12 maternal families with 20 seeds per family from each population (for a full description of data collection methods, see Supplementary Methods). Following a 6- week stratification at 4°C (Willemsen, 1975), we placed seeds in a 30°C germination chamber with 12h light/dark cycle. Two weeks after germination, we planted a randomly selected seedling from each maternal line in a random order into 100ml kwikpot trays with Debco Seed Raising Superior Germinating Mix. We top-watered all plants and artificially manipulated daylight following the light cycle at 47.3°N (median latitude over all sampling locations). One month later, we performed a second transplant (hereafter day 0) to 0.7L pots with Debco mix and 1.5ml slow-release fertilizer (Osmocote Pro, eight to nine months). We examined variation in life-history traits including growth, phenology and vegetative and reproductive allocation (Table 1).

**Table 1.** Traits included in this study.

| Trait | Description |
| --- | --- |
| Max. height | Maximum measured height |
| Total biomass | Above- and belowground dry biomass |
| Max. growth rate | Sensu Chuine *et al.* (2001) |
| Flowering onset | First recorded day of flowering (number of days after second transplant); first day of pollen release (male function) or receptive female function |
| Dichogamy | First recorded day of pollen release - first recorded day of receptive female function (a positive value is protogyny, a negative value is protandry) |
| Floral sex allocation (female/male) | Female (seeds) / male (raceme) dry weight (a value >1 is higher biomass allocation to female function) |
| Weight per seed | Dry weight per seed in milligrams, averaged over 20 seeds (where available) |
| Total reproductive biomass | Female (seeds) and male (raceme) dry weight |
| Relative reproductive biomass | Total reproductive biomass / total plant biomass |
| Specific leaf area | Leaf area of fully expanded fresh leaf/leaf dry weight |

To assess neutral genetic differentiation underlying trait variation resulting from non-adaptive evolutionary processes, we extracted DNA from leaf tissue of 861 individuals and performed double-digest genotype-by-sequencing library preparation (see Supplementary Methods). We aligned and filtered raw sequences following van Boheemen *et al.* (2017). Briefly, SNPs were aligned using BWA-mem (Li & Durbin, 2009) to a draft reference genome for *A. artemisiifolia* (van Boheemen *et al.*, 2017). We called variants with GATK HaplotypeCaller and filtered SNPs using GATK hard-filtering recommendations (McKenna *et al.*, 2010; Van der Auwera *et al.*, 2013). We identified a total of 11,598 polymorphic biallelic SNPs with 50% SNP call rate. We inferred population genetic structure and calculated individual and population level q-scores for the most likely number of clusters K (=2) with the Bayesian clustering method STRUCTURE v2.3.4 (Pritchard *et al.*, 2000). We used these STRUCTURE q-values as a measure of genetic and, therefore, trait differentiation resulting from non-adaptive (neutral) mechanisms.

### Statistical analyses

We conducted all statistical analyses in R v3.4.3 (R Core Team, 2017). We improved normality and reduced heteroscedacity of the data by square root or log-transforming traits where appropriate. We tested all univariate linear mixed models using the lme4 package (Bates *et al.*, 2014). All models tested responses of the following traits: maximum plant height, total biomass, maximum growth rate, flowering onset, dichogamy, floral sex allocation, weight per seed, total and relative reproductive biomass and specific leaf area (Table 1).

To explore patterns of latitudinal trait divergence among ranges, potentially indicative of local adaptation, we tested population mean trait responses to range (native North America, introduced Europe and Australia), latitude, their interaction and latitude^2^ (to account for non-linear trait responses) in multi- and univariate models ((M)ANCOVA). In these analyses, latitude values are not randomly distributed among ranges due to the geographic distribution of *A. artemisiifolia*, suggesting a violation of independence (Miller & Chapman, 2001). However, the values of the covariate (latitude) are observational and not manipulated by the independent variables (range) and the (M)ANCOVA assumptions are thus not violated (Keppel, 1991). We increased the power of the multivariate analysis (Scheiner, 2001) by removing highly correlated traits (Spearman’s ρ^2^>0.6, Table S3, Supplementary Material) and calculated the approximate F-statistics and Wilks’ λ (multivariate F-value) to measure the strength of the associations. To account for demographic history in patterns of trait divergence in the univariate mixed models, we included population mean STRUCTURE q-scores as a random effect. Here, we calculated significance of fixed effects using type III Wald F tests with Kenward-Roger’s approximation of denominator degrees of freedom and step-wise removed non-significant (p>0.05) fixed effects using the lmerTest package (Kuznetsova *et al.*, 2015), starting with the highest order interactions. To reduce false discovery of associations due to the number of traits being tested, we ‘fdr’ corrected p-values (hereafter q-values) of the combined final models using the p.adjust function, further reducing models when applicable.

To explore differences in latitudinal trait clines between ranges as revealed in the ANCOVAs, we tested for significant two-way interactions between range and latitude which is indicative of non-parallel trait~latitude slopes among the ranges. To further dissect the extent of trait divergence and its dependence on latitude, we compared ANCOVA model estimates of traits in the introduced ranges to the native estimates at the minimum and maximum observed latitude in each range, where applicable (EU_min_, NA_max_, NA_min_ and AU_max_) (Fig. S4, Supplementary Material). We tested overall pairwise range differences in trait values for significant range effects, when higher order interactions involving range were not significant. We explored the highest order significant interactions, using χ^2^ tests with Holm p-value correction using the phia package (De Rosario-Martinez, 2013).

We verified the presence of the well-described plant height-flowering time trade-off and examined associated patterns in other reproductive traits (dichogamy and sex allocation) within ranges. We tested linear relations between individual trait values of control treated plants in mixed models, with plant height, range and their interaction as fixed effects. In addition to individual STRUCTURE q-values, we added population as random effect. As above, we calculated fixed effects significance with type III Wald F tests, step-wise removing non-significant fixed effects. A significant interaction between range and height indicated a differential slope between the focal reproductive trait and plant height between ranges. We explored the highest order significant interactions using χ^2^ tests with Holm p-value correction using the phia package (De Rosario-Martinez, 2013).

To explore the impact of heterozygosity on fitness related traits we calculated heterozygosity (H_O_) as the proportion of heterozygous loci out of the total number of called genotypes for each individual. Introductions could diminish or increase heterozygosity, which in turn could inhibit or stimulate invasion (e.g. inbreeding due to genetic drift or heterosis following admixture). First, we investigated geographical patterns in genetic diversity of populations by testing the effect of range, latitude, their interaction and latitude^2^ on population mean H_O_. To explore the effect of heterozygosity on selected traits, we included population mean H_O_, latitude, their interactions with range and latitude^2^ as fixed effects. Both analyses included population STRUCTURE q-values as a random effect. Again, we calculated fixed effects significance with type III Wald F tests, step-wise removing non-significant fixed effects and explored the highest order significant interactions using χ^2^ tests with Holm p-value correction with the phia package (De Rosario-Martinez, 2013). To reduce false positive association due to multiple testing, we only tested the response of growth (plant height and biomass) and fitness (total reproductive biomass and average seed weight) related traits. As we identified signatures of repeated local adaptation in floral sex allocation (Results), we also tested the response of this trait. To reduce false discovery of associations due to the number of traits being tested, we ‘fdr’ corrected p-values (hereafter q-values) of the combined final models using the p.adjust function, further reducing models when applicable.

## RESULTS

### Climate niche dynamics

Niche overlap (*D*) was significantly lower than expected between the North American native and Eurasian invasive range (*P*<0.001) although the native and Australasian range did not have significantly lower overlap than expected (*P*=0.425). However, niche stability was low between the native and both invasive ranges (*P<*0.001). Climatic niche unfilling and expansion were not significantly different than the null model, except that especially low expansion was found in the Eurasian population (*P*=0.017), meaning that while the niche shifted, the species did not tend to occupy completely novel climates in its Eurasian range. In both introduced ranges the niche centroid shifted significantly towards hotter, wetter climates (*P*<0.001; Fig. 1). We note that this climate analysis compares the entire Eurasian and Australasian ranges with the native North American range.

**Figure 1.**
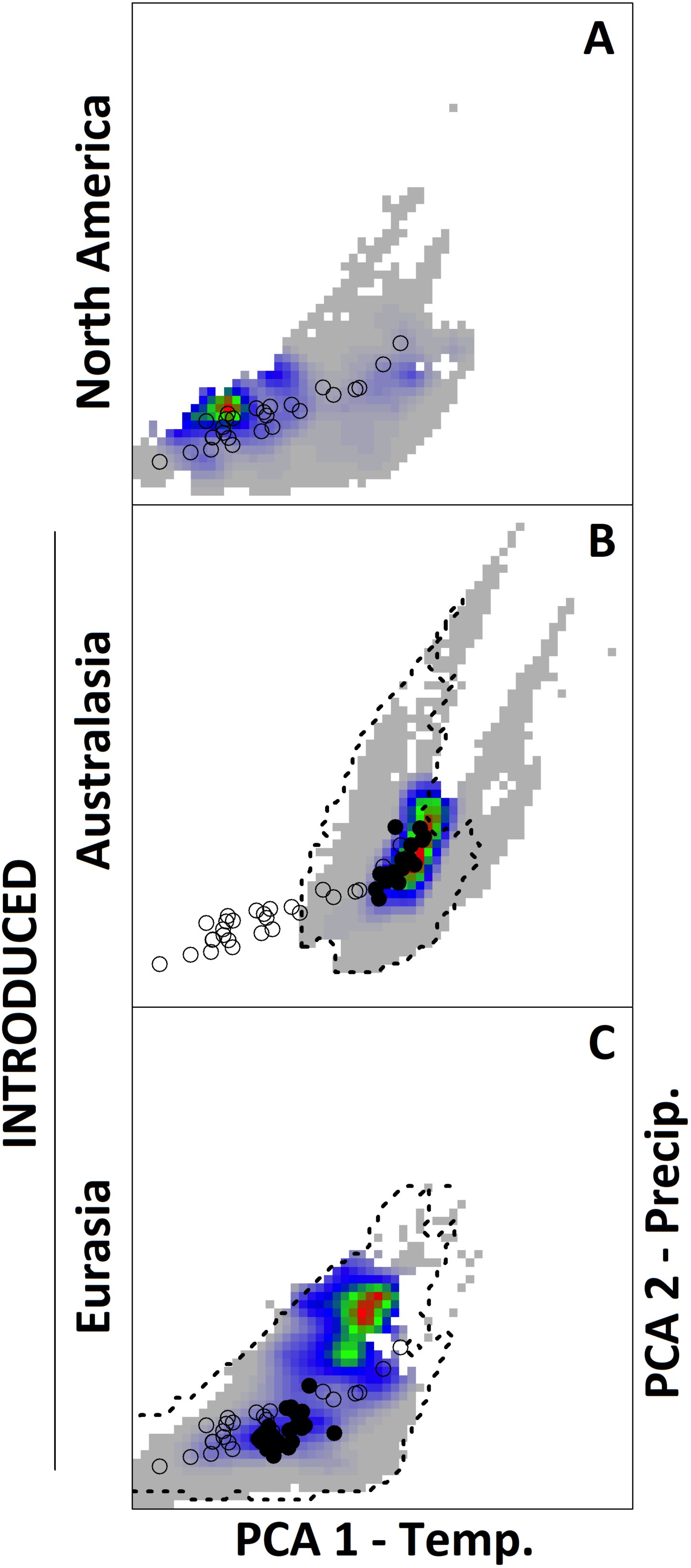
Climatic niche models of the native North American population (A) and introduced Australasian (B) and Eurasian (C) populations. Each panel shows the climate space occupied in the respective range, with a PCA variable corresponding to temperature on the x-axis and a PCA variable corresponding to precipitation on the y-axis. Color indicates occurrence probability in a given climate (red: high occurrence, green: medium occurrence, blue: low occurrence, grey: no occurrence). Open circles plot climates of the North American source localities. Closed circles plot the climates of the respective introduced range. On panels B and C, the dashed line encloses the climates shared by both the native North American and respective introduced range.

### Climate of sampled populations

For our common garden analysis we focused only on specific latitudinal transects in Europe and Australia to examine how traits have evolved along latitudinal clines during invasion, and did not include any Asian populations. We include this more general analysis of climate niche to assess how climate shifts might contribute to trait divergence among the ranges generally, although our sampling for the common garden was more limited. Climatic variables across all populations in the native North American, introduced European and introduced Australian ranges could be effectively summarized by the first two principle components (PC) in the PCA (Fig. S2, Table S2, Supplementary Material), which together explained 70.06% of between-population variation. Here, PC1 was strongly associated with latitude, temperature and seasonality, whereas PC2 was mostly precipitation related (Fig S2, Table S2). The climate experienced by Australian populations was distinct from the North American and European ranges (Fig. S2, Table S2), with higher annual, winter and summer temperatures, and with lower seasonal variation. Moreover, given the sub-tropical location of Australian populations, the sampled populations experienced higher precipitation during the growing season (Fig. S2).

### Repeatability in trait clines among ranges

Traits across all North American, European and Australian ranges were well summarized by the PCA, where the first two PCs explained 83.1% of all variation (Fig. S3, Supplementary Material). The main PCs were associated with each trait to a similar extent so no trait syndromes were apparent. Traits measured in Australian plants were generally distinct from the other populations although there was some overlap in multivariate space among the ranges (Fig. S3, Table S3). Multivariate trait analyses revealed a significant two-way interaction between latitude and range (F_14,120_=1.796, p=0.047, Wilks’ λ=0.684) (Table S4) suggesting latitudinal trait clines exist, but do not have the same relationship within ranges for all traits. Further dissection of these patterns in univariate analyses revealed maximum growth rate, flowering onset, dichogamy, average seed weight, total reproductive biomass and specific leaf area (SLA) displayed similar latitudinal clines among ranges, indicated by a significant latitude effect but an absence of a higher-order interaction (Fig. 2, Table 2). We identified range differences in latitudinal trait clines for maximum height, total biomass, floral sex allocation and relative reproductive biomass, as indicated by significant range:latitude interactions. However, all of these slopes were significantly different from zero and were in the same direction as the native North American patterns (Fig. 2, Table S5a).

**Figure 2.**
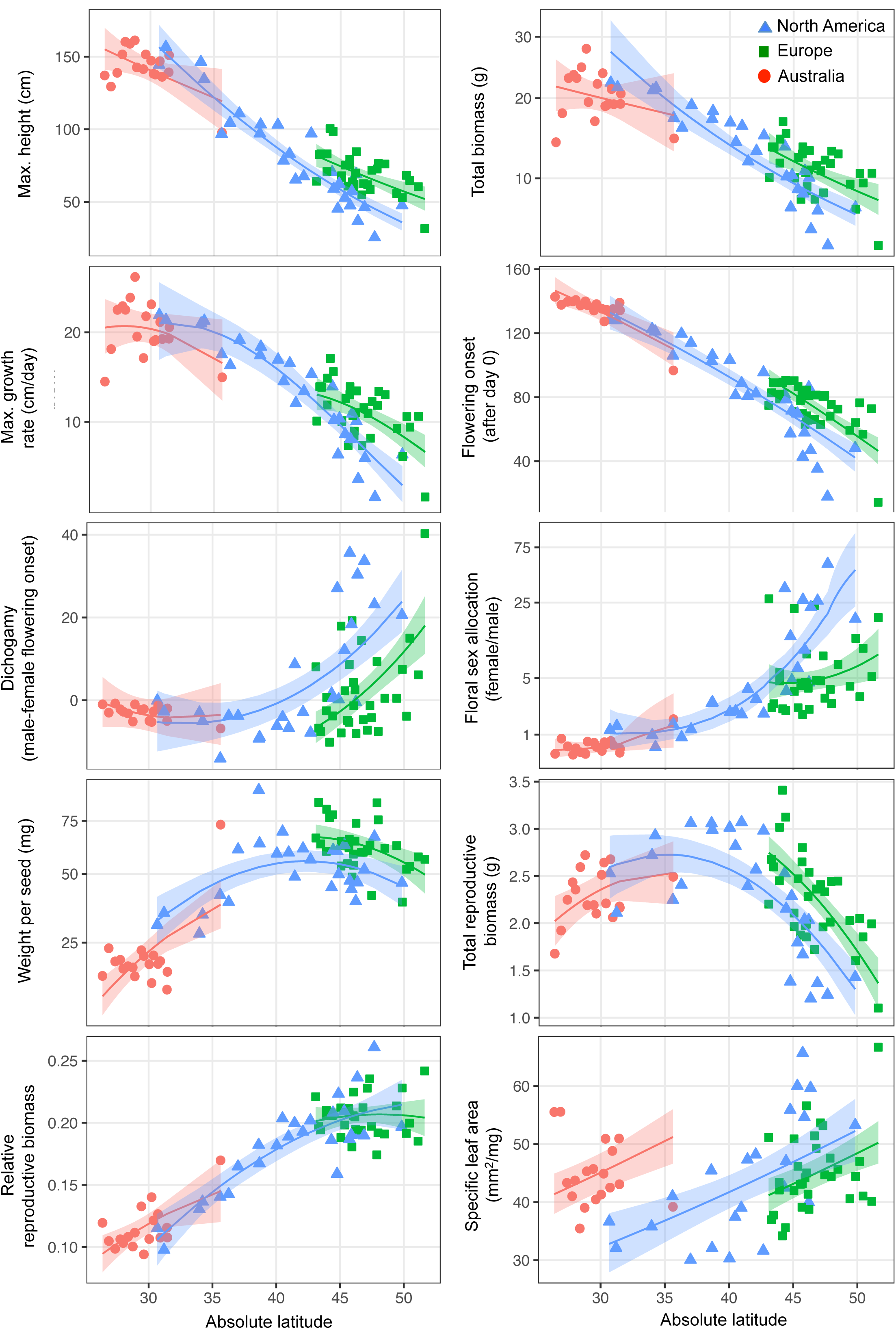
Traits responses (population means) to absolute latitude in the native North American (blue triangles), introduced European (green squares) and Australian (red circles) ranges, with model predictions and 95% shaded confidence intervals from step-wise reduced models (Table 2).

**Table 2.**
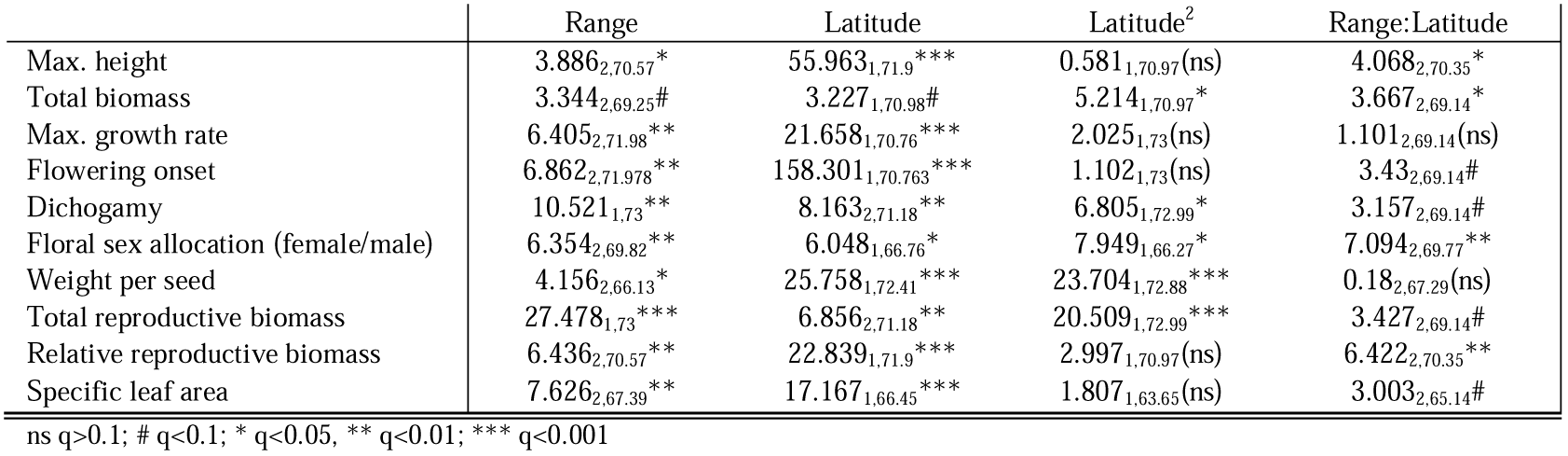
*Ambrosia artemisiifolia* population mean trait responses to range, latitude, their interaction and latitude^2^, with population q-values as random effects. We reported Wald type III F-values, with Kenward-Roger degrees of freedom as subscript and symbols specifying significance (fdr corrected q) of effect.

At higher latitudes, plants were shorter, weighed less, reached lower maximum growth rates and flowered earlier. Flowering onset extremes were 14-133 days after transplant (population means for EU20 and AU13, Fig. 2). In all ranges, dichogamy (the temporal separation of pollen dispersal and emergence of receptive stigma within an individual plant) was prevalent. Protogyny (emergence of stigma prior to pollen release) predominated at higher latitudinal populations, with receptive stigmas being visible up to 40 days before any pollen was released within the same plant (EU20)(Fig. 2). Conversely, protandry prevailed at latitudes below 40**°** from the equator, with pollen release occurring up to 14 days before stigmas became receptive (NC)(Fig. 2). Floral sex allocation followed a similar trend across ranges, with a slight male function bias at lower latitudes, shifting towards an extreme female function bias at high-latitude populations (Fig. 2). The biggest seeds were found at mid-latitudinal populations (38.6**°**N in KY), with seeds decreasing in size towards the high and low latitudes (Fig. 2, Table 2). Total reproductive biomass also showed a similar curved relationship, with the combined weight of racemes (male floral sex function) and seeds (female floral sex function) being up to three times as high at mid-latitudinal populations compared to high-latitudinal plants. In contrast, the relative reproductive biomass increased with latitude. Within each range, plants from lower latitudes had lower SLA (Fig. 2).

### Trait divergence between ranges

While latitudinal trait clines were repeatable for many traits as described above, we identified shifts in trait values at comparable latitudes as revealed by significant range effects (Table 2). Maximum growth rates were highest in Europe and lowest in Australia (Fig. 2, Table 3a). European plants also flowered later than North American and Australian plants at similar latitudes. The temporal separation between pollen release and the appearance of receptive stigma (dichogamy) was greater in the native North America compared to Europe (Fig. 2, Table 3a). European seeds were heavier and plants had higher total reproductive biomass than those measured in the other ranges. At any given latitude, Australian leaves had higher SLA compared to the native range, with lowest SLA in European populations (Fig. 1, Table 3a).

**Table 3:** Range differences of *A. artemisiifolia* population mean traits at comparable latitudes for significant (q<0.05) range effects (a, Table 2) and trait differences between ranges at minimum (min) and maximum (max) latitudes (Figure S4, Supplementary Material) for significant range:latitude interactions by comparing trait values at (b, Table 2).

| a) | North America -<br>Europe | North America -<br>Australia | Europe -<br>Australia |
| --- | --- | --- | --- |
| Max. growth rate | 12.328<sub>1</sub>** | 4.264<sub>1</sub>* | 1.278<sub>1</sub>** |
| Flowering onset | 16.465<sub>1</sub>*** | 0.123<sub>1</sub>(ns) | 13.269<sub>1</sub>*** |
| Dichogamy | 18.8<sub>1</sub>*** | 0.077<sub>1</sub>(ns) | 4.87<sub>1</sub># |
| Weight per seed | 7.44<sub>1</sub>* | 2.059<sub>1</sub>(ns) | 6.002<sub>1</sub>* |
| Total reproductive biomass | 15.228<sub>1</sub>*** | 0.811<sub>1</sub>(ns) | 6.516<sub>1</sub>* |
| Specific leaf area | 2.734<sub>1</sub># | 14.936<sub>1</sub>*** | 14.736<sub>1</sub>*** |

| ) | North<br>America-Europe | North<br>America-Australia | Europe-Australia |
| --- | --- | --- | --- |
| Max. growth rate | 13.269<sub>1</sub>*** | 4.178<sub>1</sub>* | 11.606<sub>1</sub>** |
| Flowering onset | 16.465<sub>1</sub>*** | 0.123<sub>1</sub>(ns) | 13.269<sub>1</sub>*** |
| Dichogamy | 18.8<sub>1</sub>*** | 0.077<sub>1</sub>(ns) | 4.87<sub>1</sub># |
| Weight per seed | 7.44<sub>1</sub>* | 2.059<sub>1</sub>(ns) | 6.002<sub>1</sub>* |
| Total reproductive<br>biomass | 15.228<sub>1</sub>*** | 0.811<sub>1</sub>(ns) | 6.516<sub>1</sub>* |
| Specific leaf area | 2.734<sub>1</sub># | 14.936<sub>1</sub>*** | 14.736<sub>1</sub>*** |

| b) | North America (NA) - Europe (EU) |  | North America (NA) - Australia (AU) |  |
| --- | --- | --- | --- | --- |
| Trait | EU<sub>min</sub> | NA<sub>max</sub> | NA<sub>min</sub> | AU<sub>max</sub> |
| Max. height | 8.679<sub>1</sub>** | 25.236<sub>1</sub>*** | 4.818<sub>1</sub># | 0.039<sub>1</sub>(ns) |
| Total biomass | 0.339<sub>1</sub>(ns) | 15.572<sub>1</sub>*** | 0.564<sub>1</sub>(ns) | 1.307<sub>1</sub>(ns) |
| Floral sex allocation (female/male) | 0.012<sub>1</sub>(ns) | 26.417<sub>1</sub>*** | 1.586<sub>1</sub>(ns) | 0.096<sub>1</sub>(ns) |
| Relative reproductive biomass | 4.289<sub>1</sub>(ns) | 7.557<sub>1</sub>* | 0.065<sub>1</sub>(ns) | 0.017<sub>1</sub>(ns) |

ns $q > 0.1$ ; # $q < 0.1$ ; \* $q < 0.05$ , \*\* $q < 0.01$ ; \*\*\* $q < 0.001$

Dissection of range differences in latitudinal trait clines (maximum plant height, total biomass, floral sex allocation and relative reproductive biomass) revealed most significant interactions between range and latitude were prompted by clinal differences between the introduced European and native North American populations (Table S5, Supplementary Material). For these traits, the discrepancy between North American and European trait values increased with increasing latitudes, such that at high-latitude populations, European plants were taller, heavier and less female-biased in floral sex allocation (Table 3b). Moreover, Australian plants found closest to the equator were significantly shorter than native North American expectations (Table 3b).

### Trade-offs between life-history traits

We tested for the presence of a trade-off between plant height and flowering time and investigated associated patterns in dichogamy and floral sex allocation and height. As expected, taller plants started flowering later in both the native and the introduced European range, though this pattern was not significant in introduced Australian populations (Fig. 3, Table 4, Table S6, Supplementary Material). Correspondingly, we observed protogyny and a large female-biased sex allocation in short plants versus protandry with a slight male bias in tall plants. These dichogamy associations with height were not significant in Australia (Fig. 3, Table 4 & S6). However, it is possible that the height of some large Australian plants might have been truncated due to greenhouse conditions.

**Figure 3.**
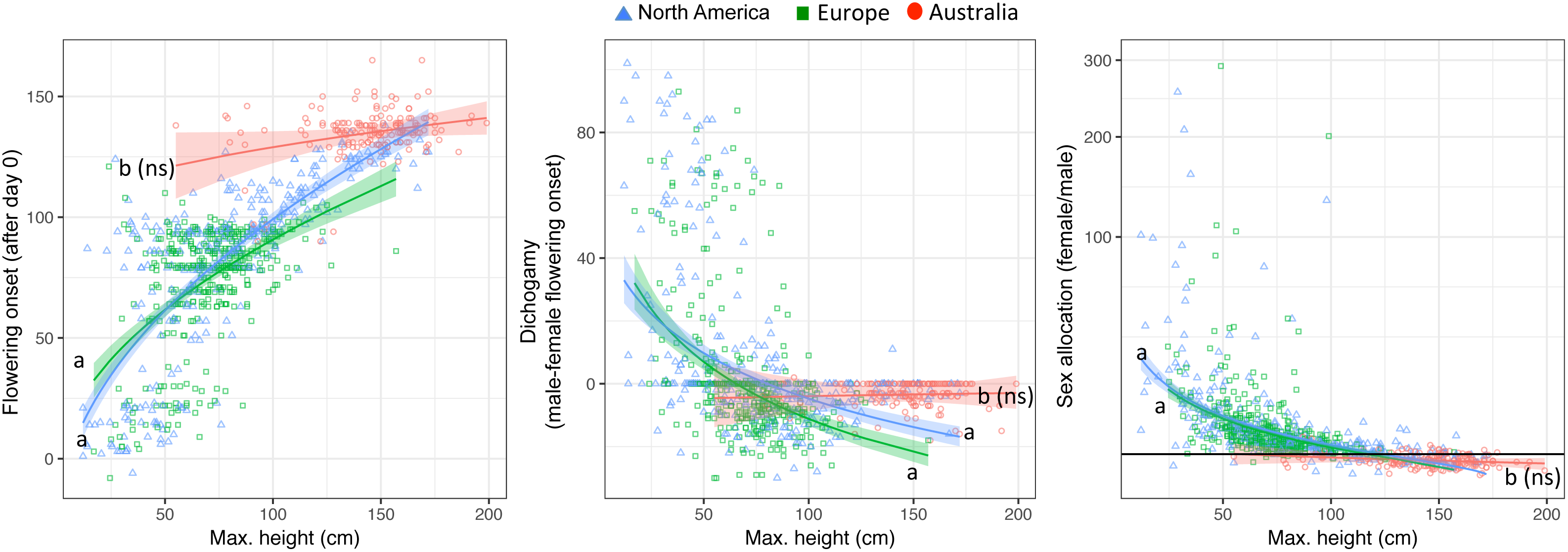
Flowering time, dichogamy and floral sex allocation responses to maximum plant height (individual values) in the native North American (NA, blue triangles), introduced European (EU, green squares) and Australian (AU, red circles) ranges, with model predictions and 95% shaded confidence intervals from step-wise reduced models (Table 4). Differences in slopes are indicated by letters and are significantly different from zero unless otherwise indicated (ns)(Table S5, supporting information).

**Table 4:** Flowering time and sex function allocation responses to maximum plant height (Height), range and their interaction, with individual q-values and population as random effects. We reported Wald type III F-values test, Kenward-Roger degrees of freedom as subscript and symbols specifying significance of effect.

| Trait | Range | Height | Range:Height |
| --- | --- | --- | --- |
| Flowering onset | 20.658<sub>2,606.14</sub>*** | 105.082<sub>1,766.54</sub>*** | 14.767<sub>2,671.02</sub>*** |
| Dichogamy | 6.503<sub>510.44</sub>** | 51.235<sub>1,670.96</sub>*** | 10.621<sub>2,545.18</sub>*** |
| Floral sex allocation (female/male) | 17.451<sub>2,508.47</sub>*** | 138.084<sub>1,659.31</sub>*** | 16.963<sub>2,508.47</sub>*** |

ns q>0.1; # q<0.1; \* q<0.05, \*\* q<0.01; \*\*\* q<0.001

### Associations between heterozygosity and life-history traits

To identify geographic patterns in observed heterozygosity (H_O_), we tested the effect of range, latitude and their interaction on H_O_. We found no latitudinal patterns in H_O_ varying within ranges (range:latitude, χ^2^_1_=3.811, p=0.149) or among all ranges (latitude, χ^2^_1_=0.000, p=0.986). We did identify variable H_O_ between ranges (range, (χ^2^_1_=6.446, p=0.040), resulting from significantly lower H_O_ in Australia compared to native North America (χ^2^_1_ =6.446; p=0.033). When accounting for latitude and population genetic structure, we found a significant interaction effect between mean population H_O_ and range on total biomass (Fig. 4, Table 5). Pairwise range comparisons in post-hoc tests revealed a higher H_O_ that was associated with heavier Australian plants (Table S7, Supplementary Material). Moreover, we found that mean population H_O_ was positively correlated with seed size in all ranges (Fig. 4, Table 5). We found no effect of individual genomic heterozygosity on plant height, phenology, dichogamy, total or relative reproductive investment and floral sex allocation (Table 5).

**Figure 4.**
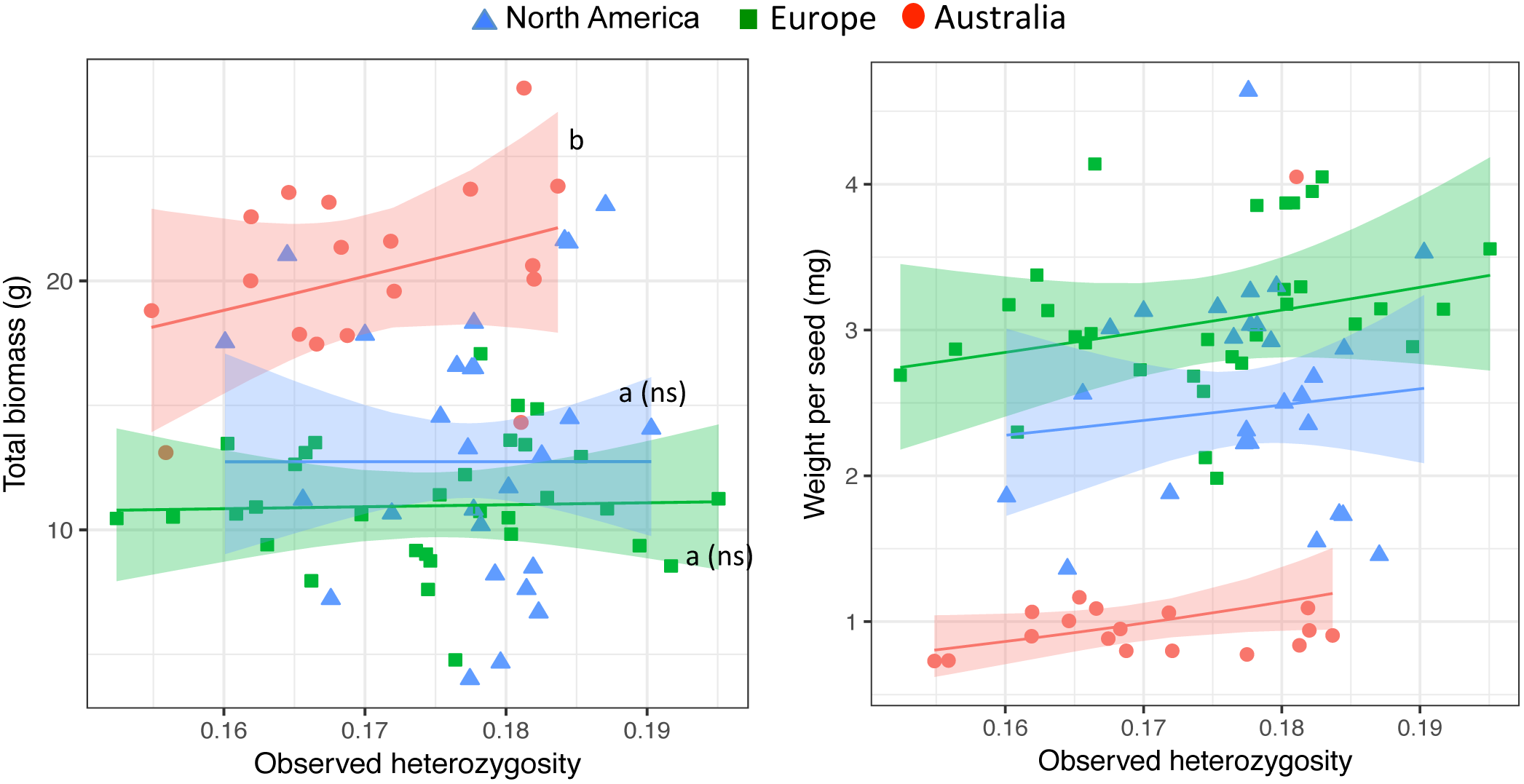
Total biomass and weight per seed response to heterozygosity (population means) in the native North American (blue triangles), introduced European (green squares) and Australian (red circles) ranges, with model predictions and 95% shaded confidence intervals from step-wise reduced models (Table 5). Slopes of predicted lines are significantly different from zero, unless otherwise indicated as (ns).

**Table 5:**
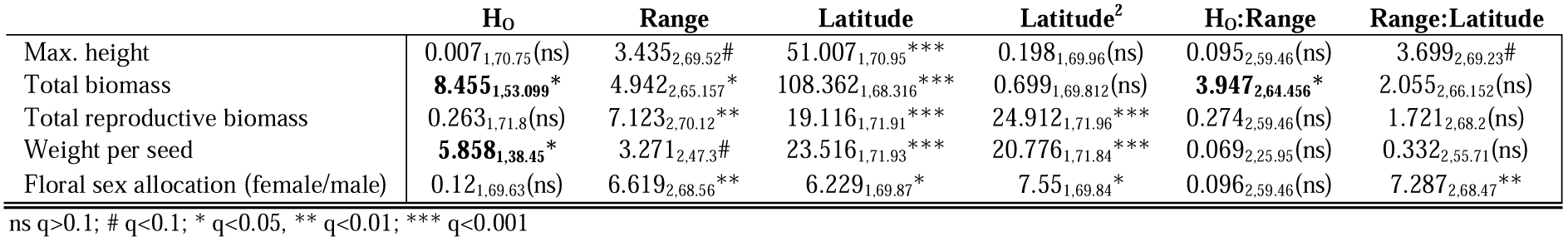
Trait responses (population mean) to range, latitude, latitude^2^, heterozygosity (H_O_) and interactions, with population q-values as random effects. We reported Wald type III F-values test values, Kenward-Roger degrees of freedom as subscript and symbols specifying significance of effect.

## DISCUSSION

We show genetically based differentiation along multiple latitudinal clines in all examined traits including plant size, growth, reproductive investment, phenology, dichogamy, SLA, and sex allocation. Remarkably, the clinal patterns apparent in the native range evolved repeatedly within both introduced ranges over the course of only 100-150 years and despite limited neutral genetic variation in the introduced Australian range. These patterns are consistent with rapid adaptation, as we accounted for neutral genetic differentiation. Moreover, low effects of maternal environment are expected (Hodgins & Rieseberg, 2011) and the introduction history of this species (van Boheemen *et al.*, 2017) reveals climate-matching (Maron *et al.*, 2004) is unlikely. The adaptive trait divergence at similar latitudes, together with a centroid shift to warmer and wetter climates in the introduced ranges, could suggest invasive populations have adapted to more productive environments following introduction. The observed rapid evolution has implications for the evolutionary potential of this species and further range expansion following climate change. Furthermore, the divergence of reproductive traits such as flowering time, sex allocation and seed size during recent range expansion should impact the production of allergenic pollen as well as the abundance and dispersal of seed that could impact spread.

### Climate niche shifts

Higher resource levels, such as increased water availability, are a known contributor to invasion in many plant species (Blumenthal, 2006; Dlugosch *et al.*, 2015b). Increased resource availability may occur through a shift in the fundamental or realized niche during invasion. The latter can result from reductions in competition, perhaps reflecting the presence of a vacant niche in the introduced range (e.g. Dlugosch *et al.*, 2015b). Climate niche dynamics analysis reveals higher *A. artemisiifolia* abundance in warmer and wetter climates in the introduced ranges compared to the native range. It is possible that this centroid shift reflects an historic effect where colonization of warmer and wetter environments occurred earlier, or perhaps by genotypes pre-adapted to these climates (*but see* van Boheemen *et al.*, 2017). Alternatively, the shift might reflect changes in biotic interactions leading to greater abundance of this species in high resource environments or differences in the availability of these climates in the introduced regions. Evolutionary processes that allow introduced species to colonize warmer and wetter environments than those occupied by native plants could also cause centroid shifts. This evolutionary interpretation is supported as Australian populations follow trait trajectories parallel to, but extending beyond, those of the native range.

### Repeated latitudinal clines

Our common garden experiments using samples collected across multiple similar latitudinal gradients, reveal that local adaptation can happen quickly and predictably, with repeated evolution of native clines in both of the introduced ranges. Latitudinal clines in phenology and size are a common feature of many geographically widespread plant species (e.g. Colautti *et al.*, 2010; Li *et al.*, 2014), with *A. artemisiifolia* flowering shown to be driven by the association between season length and latitude (Ziska *et al.*, 2011). Short season lengths at high latitudes can select for earlier flowering (Bradshaw & Holzapfel, 2008; Colautti & Barrett, 2013), while the evolution of delayed flowering at low latitudes reflects the trade-off between size and the timing of reproductive maturity, where fitness is maximized by flowering later at a large size (Colautti *et al.*, 2010; Colautti & Barrett, 2013). This correlation between plant size and flowering time has been reported for *A. artemisiifolia* (Hodgins & Rieseberg, 2011; Leiblein-Wild & Tackenberg, 2014; Scalone *et al.*, 2016) and our results are consistent with rapid genetic differentiation in plant size, growth rates and phenology in response to latitude-associated selection pressures such as season length.

We exposed repeated patterns of genetic differentiation in sex allocation strategy over similar latitudinal clines, consistent with rapid adaptation to local climate. Plants sourced from higher latitudes displayed female-biased sex-allocation and protogyny, with more balanced floral sex allocation and a decrease in the temporal separation of male and female function towards the equator. Previous studies on *A. artemisiifolia* showed plasticity for sex-allocation and dichogamy in relation to plant size (Paquin & Aarssen, 2004; Friedman & Barrett, 2009; Friedman & Barrett, 2011) and ample genetic variation for evolution to act on (Friedman & Barrett, 2011). Local seed dispersal should lead to saturating female gain curves (Lloyd & Bawa, 1984; Sakai & Sakai, 2003), yet more linear male function gain curves are predicted in wind pollinated plants with increasing height (Klinkhamer *et al.*, 1997; Friedman & Barrett, 2009). As a result, outcrossing wind-pollinated hermaphrodites with local seed dispersal, such as *A. artemisiifolia*, are predicted to adaptively change sex allocation to be more male-biased with increase in size (Lloyd, 1984; De Jong & Klinkhamer, 1989; de Jong & Klinkhamer, 1994; Klinkhamer *et al.*, 1997), consistent with patterns observed in this study. Spatial heterogeneity has been observed in animal pollinated plants, where small, resource limited individuals often allocate more resources to male function (Korpelainen, 1998; Guo *et al.*, 2010). However, these studies are on wild populations and cannot separate environmental and genetic effects in allocation patterns along resource gradients. Our current findings from common garden experiments are therefore novel in identifying genetic differentiation among populations in sex allocation over spatial gradients.

### Trait divergence among the ranges

Many hypotheses aim to explain the success of invasive species, including the evolution of increased competitive ability (EICA) through escape from native herbivores (Blossey & Notzold, 1995). Moreover, if trade-offs between performance and abiotic stress tolerance occur, greater resources could facilitate the evolution of more competitive phenotypes in introduced ranges (Grime, 1977; Bossdorf *et al.*, 2005; He *et al.*, 2010; Dlugosch *et al.*, 2015b). Our reported trait shifts in European population compared to natives at equivalent latitudes indeed suggest an increase in competitive ability, commonly measured as elevated plant growth and reproductive effort (Felker□Quinn *et al.*, 2013). These observations might reflect the warmer and wetter European climate (Fig. 1), as no general evidence for EICA has been found in Europe (Hodgins & Rieseberg, 2011; van Boheemen *et al.*, 2018) despite shifts in herbivore community composition in Europe and Australia (Genton *et al.*, 2005; Palmer & McFadyen, 2012; Essl *et al.*, 2015). Notably, although Europe was identified as the introduction source for Australian populations (van Boheemen *et al.*, 2017), traits measured within each range were highly dissimilar. Most of the sampled Australian populations extended beyond absolute latitudes of the other populations and were situated in warmer, less seasonal climates (Fig. S2). These factors might explain Australian trait variation beyond values observed in source populations.

### Heterozygosity and invasion

Genetic drift within small founding populations and on the invasion front can lead to reduced genetic diversity, potentially impacting additive genetic variation (Dlugosch & Parker, 2008a; Excoffier & Ray, 2008; Peischl *et al.*, 2013; Bock *et al.*, 2015). In *A. artemisiifolia*, Australian populations were bottlenecked and likely subjected to high genetic drift, whereas multiple introductions into Europe from distinct native sources has implicated admixture as a driver of invasion success (van Boheemen *et al.*, 2017). We found the biomass of Australian, but not European, plants was indeed associated with heterozygosity, providing only partial support for the fitness benefits of heterozygosity during invasion (Peischl & Excoffier, 2015). Admixture and heterosis are unlikely to be main drivers of invasiveness in Europe, as we found no relationship between population level heterozygosity and any trait other than seed size. Indeed, in most experimentally admixed European and native *A. artemisiifolia* crosses heterosis was absent (Hahn & Rieseberg, 2017). These observations suggest demographic processes had very limited consequences (negative or positive) on the fitness of these introduced populations. However, local adaptation of life history traits such as plant size across broad environmental gradients may mask heterozygosity-fitness correlations. Future studies could address this question by examining the association between heterozygosity and fitness in single populations (e.g. Conte *et al.*, 2017).

In plants, reduced seed size is one trait that could aid dispersal and might therefore be expected to evolve during range expansion (Bartle *et al.*, 2013; Huang *et al.*, 2015). Spatial sorting for dispersal traits at the expansion front has been well documented in other invasions, such as the cane toads (Estoup *et al.*, 2004; Phillips *et al.*, 2006). In Europe, spatial sorting for increased dispersal, and therefore smaller seeds, could have occurred at the range edge during expansion northwards. However, this mechanism would only explain the seed size decline in low-latitudinal populations in Australia, where range expansion likely occurred both north- and southward (Palmer & McFadyen, 2012; van Boheemen *et al.*, 2017). Moreover, recent evidence shows dispersal distance is determined to a much larger extent by plant height than seed traits (Thomson *et al.*, 2011; Tamme *et al.*, 2014; Augspurger *et al.*, 2017). The association between seed size and mean population heterozygosity we identified in all three ranges could be expected when small seeds aid dispersal, as founder effects should also reduce heterozygosity during colonization. Though additional factors likely shape seed size evolution, our findings suggest seed size divergence could represent an important difference in life-history strategies between ranges. Moreover, we observed patterns indicating a relationship between genomic and ecological dynamics potentially linked to range expansion and colonization.

### Conclusion

Invasive species often exhibit rapid adaptation despite facing novel selective pressures (Lachmuth *et al.*, 2011; Colautti & Barrett, 2013; Chown *et al.*, 2014; Turner *et al.*, 2014). Moreover, the success of invasives is considered paradoxical as strong demographic changes are predicted to enhance inbreeding and reduce genetic variation and, consequently, evolutionary potential (Allendorf & Lundquist, 2003).

We investigated these apparent contradictions in a comprehensive study. We compared the native range with multiple introduced ranges with highly distinct demographic histories, characterized similarities and shifts in climatic niches, tested adaptation in a large number of life-history traits and explored heterozygosity-fitness associations while accounting for non-adaptive population differentiation. We found strong evidence for parallel adaptation in all three ranges. This study therefore emphasizes that although introduction dynamics can affect genetic diversity (Dlugosch & Parker, 2008a) the adaptive potential of those traits might not be constrained to a similar extent (Dlugosch *et al.*, 2015a).

## ACKNOWLEDGEMENTS

We would like to J. Vamosi and three anonymous reviewers for their invaluable comments. We thank J. Stephens and A. Wetherhill for sample collection, S. Bou-Assi, M. Kourtidou, J. Taylor, G. Boinnard, T. Freeman and E. Barnett for greenhouse assistance, S. Bou-Assi and K. Nurkowski for genomic analyses and T. Connallon for manuscript suggestions. A Monash University Dean’s International Postgraduate Research Scholarship was provided to LAB, a Monash University Start-up Grant to KAH.

## AUTHOR CONTRIBUTIONS

LAB and KH designed the project, with data collection and analyses carried out by LAB and refined by KH. DZA developed and carried out the niche distribution modelling. All authors discussed the results and contributed to the MS writing.

## DATA ACCESSBILITY

Sequence data are available at the National Center for Biotechnology Information (NCBI) Sequence Read Archive under Bioproject PRJNA449949.

## THE FOLLOWING SUPPORTING INFORMATION IS AVAILABLE FOR THIS ARTICLE

**Fig. S1** Sampling locations

**Fig. S2** Principle component analysis of climatic and geographic variables

**Fig. S3** Principle component analysis of traits

**Fig. S4** Graphical representation of performed trait comparisons at range edges

**Table S1** List of climatic and geographic variables

**Table S2** Correlation and principle component loadings of climatic and geographic variables

**Table S3** Trait correlation and principle component loadings

**Table S4** MANCOVA testing the effect of range treatment and latitude on selected traits

**Table S5** Trait-latitude associations within ranges and range differences

**Table S6** Flowering onset, dichogamy and floral sex allocation associations with height within ranges and range differences

**Table S7** Range differences of heterozygosity-biomass associations

**Methods S1** Detailed methods for common garden set-up and data collection

**Notes S1** Australian trait divergence

## REFERENCES

Agrawal AA, Hastings AP, Bradburd GS, Woods EC, Züst T, Harvey JA, Bukovinszky T. 2015. Evolution of plant growth and defense in a continental introduction. The American Naturalist 186: E1–E15.

Allendorf FW, Lundquist LL. 2003. Introduction: population biology, evolution, and control of invasive species. Conservation Biology 17: 24–30.

Atwater DZ, Ervine C, Barney JN. 2018. Climatic niche shifts are common in introduced plants. Nature Ecology & Evolution 2: 34.

Augspurger CK, Franson SE, Cushman KC. 2017. Wind dispersal is predicted by tree, not diaspore, traits in comparisons of Neotropical species. Functional Ecology 31: 808–820.

Barrett SC, Hough J. 2012. Sexual dimorphism in flowering plants. Journal of Experimental Botany: 67–82.

Bartle K, Moles AT, Bonser SP. 2013. No evidence for rapid evolution of seed dispersal ability in range edge populations of the invasive species *Senecio madagascariensis*. Austral Ecology 38: 915–920.

Bates D, Maechler M, Bolker B, Walker S. 2014. lme4: Linear mixed-effects models using Eigen and S4. R package version 1: 1–23.

Blossey B, Notzold R. 1995. Evolution of increased competitive ability in invasive nonindigenous plants: a hypothesis. Journal of Ecology 83: 887–889.

Blumenthal DM. 2006. Interactions between resource availability and enemy release in plant invasion. Ecology Letters 9: 887–895.

Bock DG, Caseys C, Cousens RD, Hahn MA, Heredia SM, Hübner S, Turner KG, Whitney KD, Rieseberg L. 2015. What we still don’t know about invasion genetics. Molecular Ecology 24: 2277–2297.

Bonhomme M, Chevalet C, Servin B, Boitard S, Abdallah JM, Blott S, San Cristobal M. 2010. Detecting selection in population trees: the Lewontin and Krakauer test extended. Genetics.

Bossdorf O, Auge H, Lafuma L, Rogers WE, Siemann E, Prati D. 2005. Phenotypic and genetic differentiation between native and introduced plant populations. Oecologia 144: 1–11.

Bradshaw WE, Holzapfel CM. 2008. Genetic response to rapid climate change: it’s seasonal timing that matters. Molecular Ecology 17: 157–166.

Burd M, Allen T. 1988. Sexual allocation strategy in wind-pollinated plants. Evolution 42: 403–407.

Chapman DS, Haynes T, Beal S, Essl F, Bullock JM. 2014. Phenology predicts the native and invasive range limits of common ragweed. Global Change Biology 20: 192–202.

Charnov EL. 1982. The theory of sex allocation. Monographs in Population Biology 18: 1–355.

Chauvel B, Dessaint F, Cardinal-Legrand C, Bretagnolle F. 2006. The historical spread of *Ambrosia artemisiifolia* L. in France from herbarium records. Journal of Biogeography 33: 665–673.

Chown SL, Hodgins KA, Griffin PC, Oakeshott JG, Byrne M, Hoffmann AA. 2014. Biological invasions, climate change and genomics. Evolutionary Applications 8: 23–46.

Chuine I, Aitken SN, Ying CC. 2001. Temperature thresholds of shoot elongation in provenances of *Pinus contorta*. Canadian Journal of Forest Research 31: 1444–1455.

Chun YJ, Fumanal B, Laitung B, Bretagnolle F. 2010. Gene flow and population admixture as the primary post-invasion processes in common ragweed (*Ambrosia artemisiifolia*) populations in France. New Phytologist 185: 1100–1107.

Chun YJ, V LEC, Bretagnolle F. 2011. Adaptive divergence for a fitness-related trait among invasive *Ambrosia artemisiifolia* populations in France. Molecular Ecology 20: 1378–1388.

Colautti RI, Barrett SC. 2013. Rapid adaptation to climate facilitates range expansion of an invasive plant. Science 342: 364–366.

Colautti RI, Eckert CG, Barrett SC. 2010. Evolutionary constraints on adaptive evolution during range expansion in an invasive plant. Proceedings of the Royal Society B: Biological Sciences 277: 1799–1806.

Colautti RI, Lau JA. 2015. Contemporary evolution during invasion: evidence for differentiation, natural selection, and local adaptation. Molecular Ecology 24: 1999–2017.

Colautti RI, Maron JL, Barrett SCH. 2009. Common garden comparisons of native and introduced plant populations: latitudinal clines can obscure evolutionary inferences. Evolutionary Applications 2: 187–199.

Colomer-Ventura F, Martínez-Vilalta J, Zuccarini P, Escolà A, Armengot L, Castells E. 2015. Contemporary evolution of an invasive plant is associated with climate but not with herbivory. Functional Ecology 29: 1475–1485.

Conte GL, Hodgins KA, Yeaman S, Degner JC, Aitken SN, Rieseberg LH, Whitlock MC. 2017. Bioinformatically predicted deleterious mutations reveal complementation in the interior spruce hybrid complex. BMC Genomics 18: 970.

Cristescu ME. 2015. Genetic reconstructions of invasion history. Molecular Ecology 24: 2212–2225.

De Jong T, Klinkhamer P. 1989. Size-dependency of sex-allocation in hermaphroditic, monocarpic plants. Functional Ecology: 201–206.

de Jong TJ, Klinkhamer PG. 1994. Plant size and reproductive success through female and male function. Journal of Ecology: 399–402.

De Rosario-Martinez H 2013. Phia: post-hoc interaction analysis. R package.

de Villemereuil P, Gaggiotti OE, Mouterde M, Till-Bottraud I. 2016. Common garden experiments in the genomic era: new perspectives and opportunities. Heredity 116: 249.

Dlugosch KM, Anderson SR, Braasch J, Cang FA, Gillette HD. 2015a. The devil is in the details: genetic variation in introduced populations and its contributions to invasion. Molecular Ecology 24: 2095–2111.

Dlugosch KM, Cang FA, Barker BS, Andonian K, Swope SM, Rieseberg LH. 2015b. Evolution of invasiveness through increased resource use in a vacant niche. Nature Plants 1.

Dlugosch KM, Parker IM. 2008a. Founding events in species invasions: genetic variation, adaptive evolution, and the role of multiple introductions. Molecular Ecology 17: 431–449.

Dlugosch KM, Parker IM. 2008b. Invading populations of an ornamental shrub show rapid life history evolution despite genetic bottlenecks. Ecology Letters 11: 701–709.

Essl F, Biró K, Brandes D, Broennimann O, Bullock JM, Chapman DS, Chauvel B, Dullinger S, Fumanal B, Guisan A. 2015. Biological flora of the British Isles: *Ambrosia artemisiifolia*. Journal of Ecology 103: 1069–1098.

Estoup A, Beaumont M, Sennedot F, Moritz C, Cornuet J-M. 2004. Genetic analysis of complex demographic scenarios: spatially expanding populations of the cane toad, *Bufo marinus*. Evolution 58: 2021–2036.

Estoup A, Ravigné V, Hufbauer R, Vitalis R, Gautier M, Facon B. 2016. Is There a Genetic Paradox of Biological Invasion? Annual Review of Ecology, Evolution, and Systematics 47: 51–72.

Etterson JR, Shaw RG. 2001. Constraint to adaptive evolution in response to global warming. Science 294: 151–154.

Excoffier L, Ray N. 2008. Surfing during population expansions promotes genetic revolutions and structuration. Trends in Ecology & Evolution 23: 347–351.

Facon B, Genton BJ, Shykoff J, Jarne P, Estoup A, David P. 2006. A general eco-evolutionary framework for understanding bioinvasions. Trends in Ecology & Evolution 21: 130–135.

Felker-Quinn E, Schweitzer JA, Bailey JK. 2013. Meta-analysis reveals evolution in invasive plant species but little support for Evolution of Increased Competitive Ability (EICA). Ecology and Evolution 3: 739–751.

Franks SJ, Sim S, Weis AE. 2007. Rapid evolution of flowering time by an annual plant in response to a climate fluctuation. Proceedings of the National Academy of Sciences 104: 1278–1282.

Frenne P, Graae BJ, Rodríguez-Sánchez F, Kolb A, Chabrerie O, Decocq G, Kort H, Schrijver A, Diekmann M, Eriksson O. 2013. Latitudinal gradients as natural laboratories to infer species’ responses to temperature. Journal of Ecology 101: 784–795.

Friedman J, Barrett SC. 2009. Wind of change: new insights on the ecology and evolution of pollination and mating in wind-pollinated plants. Annals of Botany 103: 1515–1527.

Friedman J, Barrett SC. 2011. Genetic and environmental control of temporal and size-dependent sex allocation in a wind-pollinated plant. Evolution 65: 2061–2074.

Gaudeul M, Giraud T, Kiss L, Shykoff JA. 2011. Nuclear and chloroplast microsatellites show multiple introductions in the worldwide invasion history of common ragweed, *Ambrosia artemisiifolia*. PLoS One 6: e17658.

Geng Y-P, Pan X-Y, Xu C-Y, Zhang W-J, Li B, Chen J-K, Lu B-R, Song Z-P. 2007. Phenotypic plasticity rather than locally adapted ecotypes allows the invasive alligator weed to colonize a wide range of habitats. Biological Invasions 9: 245–256.

Genton BJ, Kotanen PM, Cheptou PO, Adolphe C, Shykoff JA. 2005. Enemy release but no evolutionary loss of defence in a plant invasion: an inter-continental reciprocal transplant experiment. Oecologia 146: 404–414.

Gladieux P, Giraud T, Kiss L, Genton BJ, Jonot O, Shykoff JA. 2010. Distinct invasion sources of common ragweed (*Ambrosia artemisiifolia*) in Eastern and Western Europe. Biological Invasions 13: 933–944.

Griffith TM, Watson MA. 2005. Is evolution necessary for range expansion? Manipulating reproductive timing of a weedy annual transplanted beyond its range. The American Naturalist 167: 153–164.

Grime JP. 1977. Evidence for the existence of three primary strategies in plants and its relevance to ecological and evolutionary theory. The American Naturalist 111: 1169–1194.

Guisan A, Petitpierre B, Broennimann O, Daehler C, Kueffer C. 2014. Unifying niche shift studies: insights from biological invasions. Trends in Ecology & Evolution 29: 260–269.

Guo H, Mazer SJ, Du G. 2010. Geographic variation in primary sex allocation per flower within and among 12 species of *Pedicularis* (Orobanchaceae): proportional male investment increases with elevation. American Journal of Botany 97: 1334–1341.

Hahn MA, Rieseberg LH. 2017. Genetic admixture and heterosis may enhance the invasiveness of common ragweed. Evolutionary Applications 10: 241–250.

He W-M, Thelen GC, Ridenour WM, Callaway RM. 2010. Is there a risk to living large? Large size correlates with reduced growth when stressed for knapweed populations. Biological Invasions 12: 3591–3598.

Hijmans RJ, Cameron SE, Parra JL, Jones PG, Jarvis A. 2005. Very high resolution interpolated climate surfaces for global land areas. International Journal of Climatology 25: 1965–1978.

Hodgins KA, Bock DG, Rieseberg L. 2018. Trait Evolution in Invasive Species. Annual Plant Reviews online 1: 1–37.

Hodgins KA, Rieseberg L. 2011. Genetic differentiation in life-history traits of introduced and native common ragweed (*Ambrosia artemisiifolia*) populations. Journal of Evolutionary Biology 24: 2731–2749.

Huang F, Peng S, Chen B, Liao H, Huang Q, Lin Z, Liu G. 2015. Rapid evolution of dispersal-related traits during range expansion of an invasive vine *Mikania micrantha*. Oikos 124: 1023–1030.

Huey RB, Gilchrist GW, Carlson ML, Berrigan D, Serra Ls. 2000. Rapid evolution of a geographic cline in size in an introduced fly. Science 287: 308–309.

Keller SR, Sowell DR, Neiman M, Wolfe LM, Taylor DR. 2009. Adaptation and colonization history affect the evolution of clines in two introduced species. New Phytologist 183: 678–690.

Keppel G. 1991. Design and analysis: A researcher’s handbook: Prentice-Hall, Inc.

Klinkhamer PG, De Jong TJ, Metz H. 1997. Sex and size in cosexual plants. Trends in Ecology & Evolution 12: 260–265.

Korpelainen H. 1998. Labile sex expression in plants. Biological Reviews 73: 157–180.

Kuznetsova A, Brockhoff PB, Christensen RHB. 2015. Package ‘lmerTest’. R package version 2.

Laaidi M, Laaidi K, Besancenot J-P, Thibaunod M. 2003. Ragweed in France: an invasive plant and its allergenic pollen. Annals of Allergy, Asthma & Immunology 91: 195–201.

Lachmuth S, Durka W, Schurr FM. 2011. Differentiation of reproductive and competitive ability in the invaded range of *Senecio inaequidens*: the role of genetic Allee effects, adaptive and nonadaptive evolution. New Phytologist 192: 529–541.

Lee CE. 2002. Evolutionary genetics of invasive species. Trends in Ecology and Evolution 17: 386–391.

Leiblein-Wild MC, Tackenberg O. 2014. Phenotypic variation of 38 European *Ambrosia artemisiifolia* populations measured in a common garden experiment. Biological Invasions 16: 2003–2015.

Li H, Durbin R. 2009. Fast and accurate short read alignment with Burrows-Wheeler transform. Bioinformatics 25: 1754–1760.

Li X-M, She D-Y, Zhang D-Y, Liao W-J. 2014. Life history trait differentiation and local adaptation in invasive populations of *Ambrosia artemisiifolia* in China. Oecologia 177: 669–677.

Lloyd D. 1984. Gender allocations in outcrossing cosexual plants.

Lloyd DG, Bawa KS. 1984. Modification of the gender of seed plants in varying conditions. Evolutionary Biology 17: 255–338.

Maron JL, Elmendorf SC, Vilà M. 2007. Contrasting plant physiological adaptation to climate in the native and introduced range of Hypericum perforatum. Evolution 61: 1912–1924.

Maron JL, Vilà M, Bommarco R, Elmendorf S, Beardsley P. 2004. Rapid evolution of an invasive plant. Ecological Monographs 74: 261–280.

McKenna A, Hanna M, Banks E, Sivachenko A, Cibulskis K, Kernytsky A, Garimella K, Altshuler D, Gabriel S, Daly M, et al. 2010. The Genome Analysis Toolkit: a MapReduce framework for analyzing next-generation DNA sequencing data. Genome Research 20: 1297–1303.

Miller GA, Chapman JP. 2001. Misunderstanding analysis of covariance. Journal of Abnormal Psychology 110: 40.

Moles AT, Wallis IR, Foley WJ, Warton DI, Stegen JC, Bisigato AJ, Cella-Pizarro L, Clark CJ, Cohen PS, Cornwell WK. 2011. Putting plant resistance traits on the map: a test of the idea that plants are better defended at lower latitudes. New Phytologist 191: 777–788.

Moran EV, Alexander JM. 2014. Evolutionary responses to global change: lessons from invasive species. Ecology Letters 17: 637–649.

Nakazato T, Bogonovich M, Moyle LC. 2008. Environmental factors predict adaptive phenotypic differentiation within and between two wild Andean tomatoes. Evolution: International Journal of Organic Evolution 62: 774–792.

Oduor AM, Leimu R, Kleunen M. 2016. Invasive plant species are locally adapted just as frequently and at least as strongly as native plant species. Journal of Ecology 104: 957–968.

Ordoñez JC, Van Bodegom PM, Witte JPM, Wright IJ, Reich PB, Aerts R. 2009. A global study of relationships between leaf traits, climate and soil measures of nutrient fertility. Global Ecology and Biogeography 18: 137–149.

Palmer B, McFadyen RC 2012. *Ambrosia artemisiifolia* L. - annual ragweed. Biological control of weeds in Australia. Collingwood: CSIRO, 52–59.

Paquin V, Aarssen LW. 2004. Allometric gender allocation in *Ambrosia artemisiifolia* (Asteraceae) has adaptive plasticity. American Journal of Botany 91: 430–438.

Peischl S, Dupanloup I, Kirkpatrick M, Excoffier L. 2013. On the accumulation of deleterious mutations during range expansions. Molecular Ecology 22: 5972–5982.

Peischl S, Excoffier L. 2015. Expansion load: recessive mutations and the role of standing genetic variation. Molecular Ecology 24: 2084–2094.

Phillips BL, Brown GP, Webb JK, Shine R. 2006. Invasion and the evolution of speed in toads. Nature 439: 803–803.

Poorter H, Niinemets Ü, Poorter L, Wright IJ, Villar R. 2009. Causes and consequences of variation in leaf mass per area (LMA): a meta-analysis. New Phytologist 182: 565–588.

Prentis PJ, Wilson JR, Dormontt EE, Richardson DM, Lowe AJ. 2008. Adaptive evolution in invasive species. Trends in Plant Sciences 13: 288–294.

Pritchard JK, Stephens M, Donnelly P. 2000. Inference of population structure using multilocus genotype data. Genetics 155: 945–959.

R Core Team 2017. R: a language and environment for statistical computing.

Rius M, Darling JA. 2014. How important is intraspecific genetic admixture to the success of colonising populations? Trends in Ecology & Evolution 29: 233–242.

Sakai A, Sakai S. 2003. Size-dependent ESS sex allocation in wind-pollinated cosexual plants: fecundity vs. stature effects. Journal of Theoretical Biology 222: 283–295.

Savolainen O, Lascoux M, Merila J. 2013. Ecological genomics of local adaptation. Nature Review Genetics 14: 807–820.

Sax DF, Brown JH. 2000. The paradox of invasion. Global Ecology and Biogeography 9: 363–371.

Scalone R, Lemke A, Štefanić E, Kolseth A-K, Rašić S, Andersson L. 2016. Phenological variation in *Ambrosia artemisiifolia* L. facilitates near future establishment at northern latitudes. PLoS One 11: e0166510.

Scheiner SM 2001. Multiple response variables and multi-species interactions. Design and analysis of ecological experiments. New York: Chapman & Hall, 99–133.

Schoener TW. 1968. The Anolis lizards of Bimini: resource partitioning in a complex fauna. Ecology 49: 704–726.

Smith M, Cecchi L, Skjøth C, Karrer G, Šikoparija B. 2013. Common ragweed: a threat to environmental health in Europe. Environment International 61: 115–126.

Sun Y, Brönnimann O, Roderick GK, Poltavsky A, Lommen ST, Müller-Schärer H. 2017. Climatic suitability ranking of biological control candidates: a biogeographic approach for ragweed management in Europe. Ecosphere 8: e01731.

Tamme R, Götzenberger L, Zobel M, Bullock JM, Hooftman DA, Kaasik A, Pärtel M. 2014. Predicting species’ maximum dispersal distances from simple plant traits. Ecology 95: 505–513.

Taramarcaz P, Lambelet C, Clot B, Keimer C, Hauser C. 2005. Ragweed (*Ambrosia*) progression and its health risks: will Switzerland resist this invasion? Swiss Medical Weekly 135: 538–548.

Thomson FJ, Moles AT, Auld TD, Kingsford RT. 2011. Seed dispersal distance is more strongly correlated with plant height than with seed mass. Journal of Ecology 99: 1299–1307.

Turner KG, Hufbauer RA, Rieseberg LH. 2014. Rapid evolution of an invasive weed. New Phytologist 202: 309–321.

van Boheemen LA, Bou-Assi S, Uesugi A, Hodgins KA. 2018. EICA fails as an explanation of growth and defence evolution following multiple introductions. bioRxiv DOI: https://doi.org/10.1101/435271

van Boheemen LA, Lombaert E, Nurkowski KA, Gauffre B, Rieseberg LH, Hodgins KA. 2017. Multiple introductions, admixture and bridgehead invasion characterize the introduction history of *Ambrosia artemisiifolia* in Europe and Australia. Molecular Ecology 26: 5421–5434.

Van der Auwera GA, Carneiro MO, Hartl C, Poplin R, del Angel G, Levy-Moonshine A, Jordan T, Shakir K, Roazen D, Thibault J. 2013. From FastQ data to high-confidence variant calls: the genome analysis toolkit best practices pipeline. Current protocols in bioinformatics 43: 11.10. 11–11.10. 33.

Willemsen RW. 1975. Dormancy and germination of common ragweed seeds in the field. American Journal of Botany 62: 639–643.

Zhang D-Y 2006. Evolutionarily stable reproductive investment and sex allocation in plants. In: Harder LD, Barrett RD eds. Ecology and evolution of flowers. Oxford: Oxford University Press, 41–60.

Zhang YY, Zhang DY, Barrett SC. 2010. Genetic uniformity characterizes the invasive spread of water hyacinth (*Eichhornia crassipes*), a clonal aquatic plant. Molecular Ecology 19: 1774–1786.

Ziska L, Knowlton K, Rogers C, Dalan D, Tierney N, Elder MA, Filley W, Shropshire J, Ford LB, Hedberg C. 2011. Recent warming by latitude associated with increased length of ragweed pollen season in central North America. Proceedings of the National Academy of Sciences 108: 4248–4251.

